# Are Maize Stalks Efficiently Tapered to Withstand Wind Induced Bending Stresses?

**DOI:** 10.1101/2020.01.21.914804

**Authors:** Christopher J Stubbs, Kate Seegmiller, Rajandeep S. Sekhon, Daniel J. Robertson

## Abstract

Stalk lodging (breaking of agricultural plant stalks prior to harvest) results in millions of dollars in lost revenue each year. Despite a growing body of literature on the topic of stalk lodging, the structural efficiency of maize stalks has not been investigated previously. In this study, we investigate the morphology of mature maize stalks to determine if rind tissues, which are the major load bearing component of corn stalks, are efficiently organized to withstand wind induced bending stresses that cause stalk lodging.

945 fully mature, dried commercial hybrid maize stem specimens (48 hybrids, ∼2 replicates, ∼10 samples per plot) were subjected to: (1) three-point-bending tests to measure their bending strength and (2) rind penetration tests to measure the cross-sectional morphology at each internode. The data were analyzed through an engineering optimization algorithm to determine the structural efficiency of the specimens.

Hybrids with higher average bending strengths were found to allocate rind tissue more efficiently than weaker hybrids. However, even strong hybrids were structurally suboptimal. There remains significant room for improving the structural efficiency of maize stalks. Results also indicated that stalks are morphologically organized to resist wind loading that occurs primarily above the ear. Results are applicable to selective breeding and crop management studies seeking to reduce stalk lodging rates.

**Highlight:** Maize stem morphology was investigated through an optimization algorithm to determine how efficiently their structural tissues are allocated to withstand wind induced bending stresses that cause stalk lodging.

## Introduction

Stalk lodging (permanent displacement of plants from their vertical orientation) severely reduces agronomic yields of several vital crop species including maize. Yield losses due to stalk lodging are estimated to range from 5-20% annually (Flint-Garcia *et al*., 2003; Berry *et al*., 2007). Several internal and external factors contribute to a plant’s propensity to stalk lodge. External factors include wind speed (Wen *et al*., 2019), pest damage (Echezona, 2007), and disease (Dudley, 1994; Holbert et al., 1923). Internal factors include the plant’s morphology and material properties (Esechie, 1985; Robertson *et al*., 2017; Stubbs *et al*., 2018). Despite a growing body of literature surrounding the topic of maize stalk lodging, a detailed morphological investigation of the taper of maize stalks has not been reported. The purpose of this paper is to quantify changes in diameter and rind thickness of maize stalks as a function of plant height (i.e., taper) and to determine the structural efficiency of the taper of maize stalks. This study investigates stalk taper from a purely structural standpoint and other abiotic and biotic considerations that may affect stalk morphology (i.e., taper) of maize stalks are not considered.

To determine the structural efficiency of maize stalks one must both quantify the stalk taper and define probable wind loading scenarios. An efficiently tapered stalk is defined as one in which uniform mechanical stresses are produced when the plant is subjected to probable wind loading scenarios. In other words, the shape of the stalk is optimal, meaning that loads are supported with as little tissue as possible. An inefficient taper is one in which non-uniform mechanical stresses are produced. Inefficient stalks utilize more structural tissue than is necessary in some areas and less structural tissue than is necessary in other areas to withstand the loads to which they are subjected. In other words, for inefficient stalks the amount of structural tissue could be reduced without affecting the load bearing capacity of the stalk. The structural efficiency of maize stalks is of interest because efficient stalks would theoretically have more available biomass and bioenergy to devote to grain filling as compared to inefficient stalks (i.e., efficient stalks would have a higher harvest index).

As mentioned previously, both the taper and probable wind loading scenarios must be defined to determine the structural efficiency of maize stalks. The wind load exerted on a plant stalk, known as the drag force (*D_f_*), can be approximated as (Niklas, 2000):

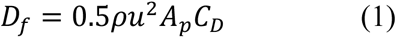

where *⍴* is the density of air, *u* is the local wind speed, *A_p_* is the projected area of the structure, and *C_D_* is the drag coefficient. While this equation appears fairly simple at first glance, it is complicated by the fact that the variables on the right hand side of the equation are functions that can vary both temporally and spatially. For example, the drag coefficient changes spatially along the length of the stalk and is also a function of the local wind speed. As the local wind speed increases, the angle of the leaf blades and tassel change (known as flagging), which alters the drag coefficient.

The strong interrelationships between the factors of Equation 1 complicate attempts to directly measure wind forces on maize stalks. Direct measurements of wind speeds have successfully been used to estimate drag forces in past studies of trees (Niklas and Spatz, 1999; Niklas, 2000). However, the large ratio of leaf area to stalk area, close proximity of maize plants to one another in commercial fields, and other confounding factors imply that a direct measurement of the wind speed near a maize stalk is not necessarily a good predictor of the drag force experienced by the stalk. Detailed computational engineering models that capture the interplay between fluid dynamics and structural deformations (i.e. fluid-structure interaction models (Zienkiewicz *et al*., 2014)) could potentially be used to calculate the drag force experienced by maize stalks over time. However, such models are computationally expensive and time-consuming to run. In summary, accurately measuring drag forces in crop canopies is challenging and remains and active area of research. An overview of this topic is given by Finnigan (Finnigan, 2000).

While direct measurement of exact wind forces on maize stalks is challenging, defining the realm of possible wind loading scenarios less so. To define the realm of possible wind loading scenarios we assume the wind speed acts in the same direction along the length of the stalk. In other words the wind does not blow in one direction at the bottom of the stalk and in a different direction at the top of the stalk. We can also bound the degree of change in the magnitude of the wind force along the length of the plant. For example, previous studies and engineering fluid mechanics theory dictate that the local wind speed in crop canopies increases with height (Cionco, 1965; Wen *et al*., 2019; Yi, 2008). A simple examination of corn stalks also suggest that the combination of the drag coefficient and projected area increases with plant height (i.e., the leaves near the bottom of mature maize plant are often dead and fall off whereas the top leaves remain structurally robust). Thus both the local wind speed and the combined effect of the drag coefficient and projected area can be assumed to increase with plant height. Combining these insights with Equation 1, we can determine that the wind force (i.e. drag force) increases with plant height. Next, we apply upper and lower bounds of probable wind loading scenarios. At the upper bound of probable wind loading we assume all of the wind force acts at the top of the plant as a point load. At the lower bound of probable wind loading we assume a uniform load is applied to stalk along its entire length (i.e., the drag force at the top of the plant is the same as the drag force in every other cross-section of the plant including the bottom of the plant). These bounds allow for all probable wind loading scenarios (e.g., linear, quadratic exponential etc. increase in drag force with plant height) and exclude improbable scenarios such as the drag force being higher at the base of the plant than at the top of the plant. Figure 1 visually represents each of these assumptions.

**Figure 1:**
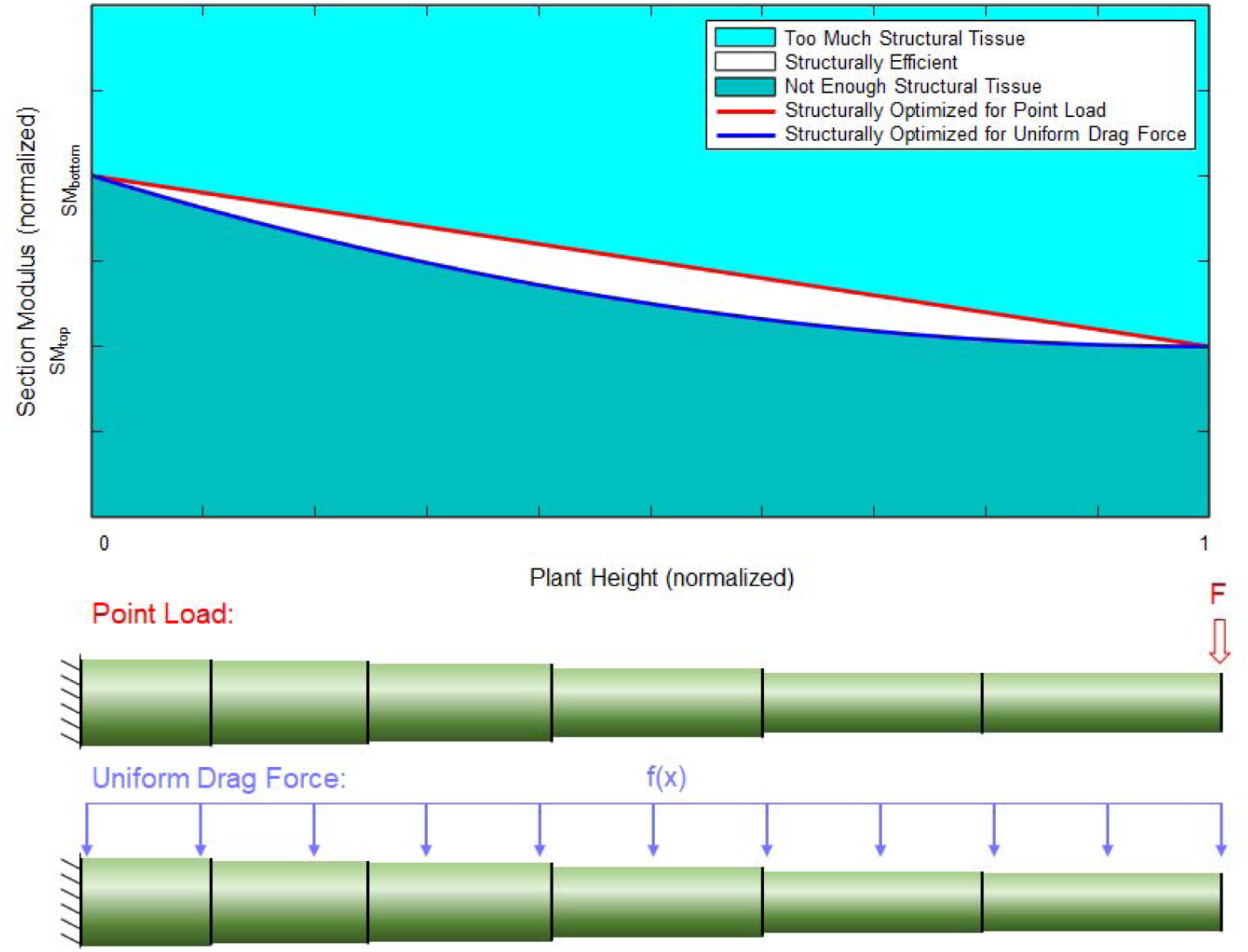
Structural efficiency along the length of the specimen as a function of probable wind loadings (top); loading diagrams for a point load and uniform drag force (bottom).

The structural efficiency of maize stalks can be determined by using Engineering equations which relate stem morphology and mechanical stress to probable wind loading scenarios presented in Figure 1. In particular, the maximum stress in any cross-section (*σ*) due to wind-induced bending is calculated as (Beer *et al*., 2002):

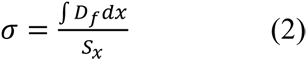

where *D_f_* is the drag force (see Equation 1) and *S_x_* is the section modulus at a distance *x* along the stalk. Section modulus is an engineering term that quantifies the morphology of the cross-section. Maize stalks possess elliptical cross-sections, therefore, the section modulus of each cross-sections is a function of the ellipse’s major diameter (*D*), minor diameter (*d*), and the thickness of the rind (*t*) in the form (Young and Budynas, 2002):

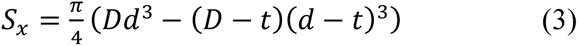

By combining Equations 2 and 3, we can calculate the drag force / section modulus that results in a uniform stress along the length of the plant.

Figure 1 displays a visual representation of the range of possible tapers for maize stalks that would produce uniform stresses during probable wind loading scenarios. The range of probable wind loadings is defined with an upper bound (red curve) of a point load applied to the top of the plant, and a lower bound (blue curve) of a uniform drag force applied to the entire length of the specimen. The graph depicts the most efficient plant tapers (white area) from the ground (x = 0) to the top of the plant specimen (x = 1), and from the section modulus at the uppermost internode of the plant specimen (y = SM_top_) to the section modulus at the bottommost internode of the plant (y = SM_bottom_). If the section modulus of an internode falls above the red curve, then that internode will experience a lower maximum stress than the rest of the plant, as it has more structural tissues (i.e., mass) than is necessary. If the section modulus of an internode falls below the blue curve, then that internode will experience a higher maximum stress than the rest of the internodes of the plant, as it has less structural tissue than is efficient. If the section modulus of an internode falls between the red and blue curves (white area), then that internode will experience a similar level of mechanical stress as compared to the rest of the plant, and therefore has an efficient allocation of structural tissues.

To determine the structural efficiency of maize plants, a select group of maize stalks were analyzed. Their major and minor diameters and rind thicknesses were measured at each internode and compared to Figure 1. In addition, a custom optimization algorithm was employed to determine the exact drag force profile for each plant that would produce the most uniform mechanical stress possible for the given stalk structure. The details and results of these experiments are presented in the following sections.

## Methods

All maize specimens in this study were subjected to the following battery of tests. First, major and minor diameters of each internode were measured with calipers and internode lengths were measured with a ruler. Second, bending strength was measured in three-point bending. Third, rind thickness was measured through rind penetration tests (Seegmiller *et al*., 2020). Each stalk was then analyzed and an optimization algorithm was employed to determine the theoretical drag force profile that would produce the most uniform stresses along the entire length of the stalk. Finally, statistical analyses were performed to investigate the relationship between each specimen’s strength and its structural efficiency.

### Plant materials

Forty-eight maize hybrids, chosen to represent a reasonable portion of maize genetic diversity, were evaluated for variation in stem morphology and structural tissue distribution. The hybrids were planted at Clemson University Simpson Research and Education Center, Pendleton, SC in well drained Cecil sandy loam soil. The hybrids were grown in a Random Complete Block Design with two replications. In each replication, each hybrid was planted in two-row plots with row length of 4.57 m and row-to-row distance of 0.76 m with a targeted planting density of 70,000 plant ha^-1^. The experiment was surrounded by non-experimental maize hybrids on all four sides to prevent any edge effects. To supplement nutrients, 56.7 kg ha^-1^ nitrogen, 86.2 kg ha^-1^ of phosphorus and 108.9 kg ha^-1^ potassium was added at the time of soil preparation, and additional 85 kg ha^-1^ nitrogen was applied 30 days after emergence. Standard agronomic practices were followed for crop management.

Stalks used for this study were harvested when all the hybrids were either at or past physiological maturity (i.e., 40 days after anthesis). Ten competitive plants from each replication were harvested by cutting at just above ground level, stripped of all the leaves and ears, and transferred to a forced air dryer for drying at 65°C. Some plots lacked 10 competitive plants and, therefore, the total number of plants evaluated for each hybrid varied slightly. In total, 945 fully mature, dried commercial hybrid maize stalks were used in this study (48 hybrids, ∼2 replicates, ∼10 samples per hybrid).

### Three-Point-Bending Tests

Specimens were tested in three-point-bending using an Instron Universal Testing System (Instron Model # 5944, Norwood, MA). Specimens were supported at their bottom and top node, and loaded at their middle node. Care was taken to ensure that the specimens were both loaded and supported at nodes, and that the span lengths were maximized. This was done to obtain the most natural possible failure modes (Robertson *et al*., 2015; Stubbs *et al*., 2018). Specimens were loaded at a rate of 2 mm/s until structural failure. Additional details on the three-point-bending test protocol were documented in a previous study (Robertson *et al*., 2017). Short span 3-pt bend tests (test of a single internode) were not employed in this study as they have been shown to produce unnatural failure patterns and result in inaccurate bending strength measurements (Robertson *et al*., 2015).

### Morphology Measurements

Internode lengths of each specimen were measured with a ruler. Other morphology measurements were taken at the midspan of each internode of every specimen. In particular, caliper measurements were used to obtain the minor and major diameters of each internode. Rind penetration tests were used to obtain the rind thickness of each internode. Rind penetration tests were performed using an Instron universal testing machine. A probe was briefly forced through the specimen at a rate of 25 mm/s, and the resulting force-displacement curve was analyzed using a custom MATLAB algorithm to calculate the rind thickness (t) of the stalk cross-section. Additional details on the rind penetration test protocol are documented in (Seegmiller *et al*., 2020).

### Optimization of Loading Condition

An optimization algorithm was employed to determine the drag force profile (f(x)) for each stem specimen that would produce the most uniform stress along the length of the stalk. As the specimens examined only spanned the bottom half of the stalk (from the ground to the ear), the loading above the ear was resolved into a single positive force (F_0_) and positive moment (M_0_) applied to the top of the specimen as described by Beer (Beer *et al*., 2002) and shown in Figure 2. For the hybrids investigated, the ear was an average of 49.6% (+/-14.3% standard deviation) of the way up the stalk.

**Figure 2:**
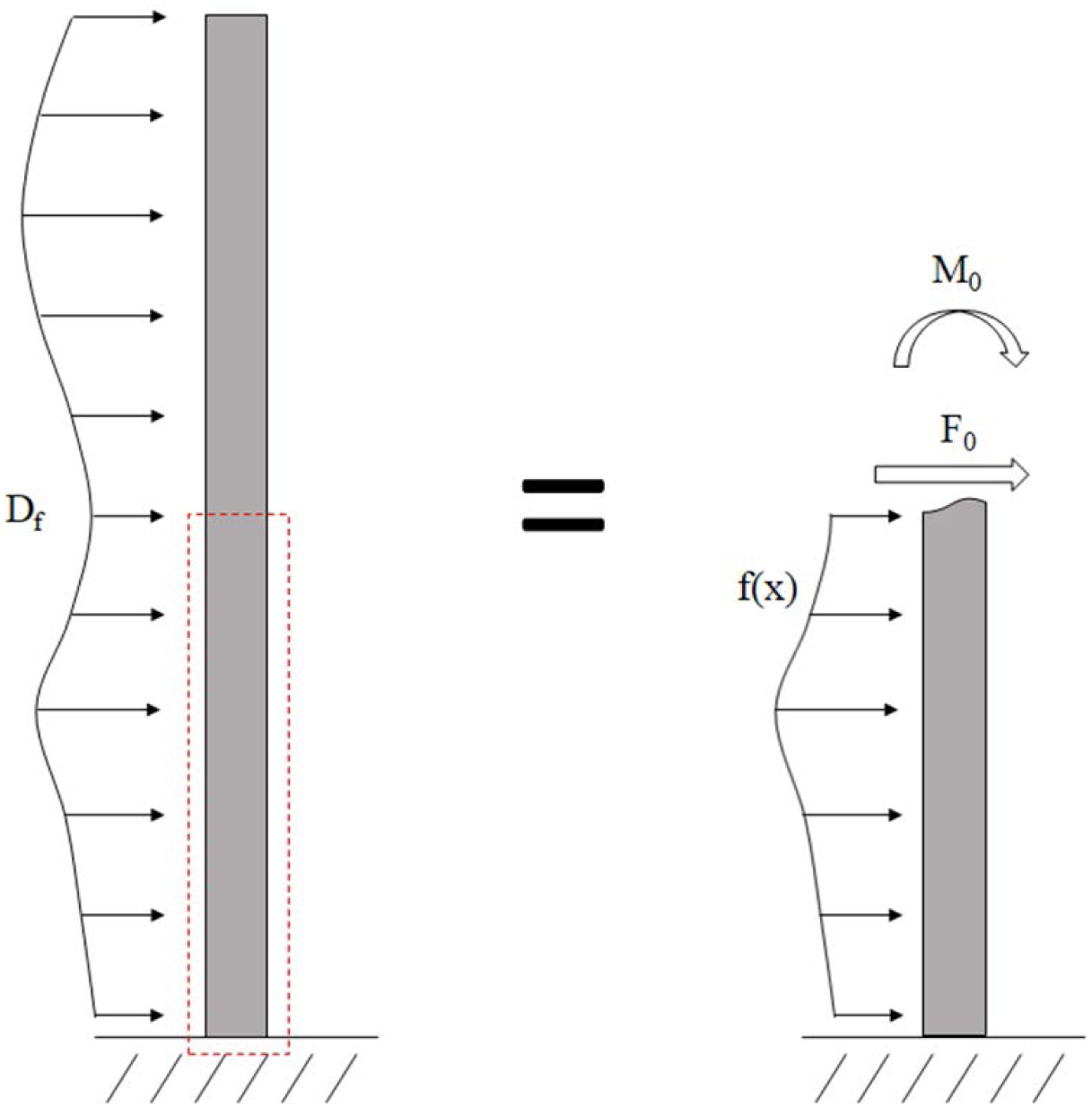
A loading diagram of the wind on the plant stalk with an unknown load distribution along the length of the stalk (left); the wind loading above the ear can be resolved as an unknown positive force (F_0_) and positive moment (M_0_) applied to the top cross section (right) (Beer et al., 2002).

Based on this setup, we can now calculate the resolved force (F_0_) and the loading profile (f(x)) for each specimen that results in the most uniform stress state in each particular specimen. This was accomplished through the use of an optimization algorithm. In particular, a custom code was developed in Matlab to perform an *fmincon* optimization for each stem specimen (Chuan et al., 2014; Han, 1977, 1977; Sreeraj et al., 2013). The objective of the optimization function was to minimize the variation in mechanical stress across the length of the specimen by changing the values of the input parameters F_0_ and f(x) (see figure 3). To accomplish this each specimen was computationally partitioned into 100 cross-sections and the optimization routine would then: (1) take in user-supplied initial estimations of F_0_ at the top of the specimen and f(x) at the other 99 cross-sections (100x1 vector ***X***), (2) calculate bending stress at every cross section using Equation 2 (100x1 vector ***s***), (3) calculate the total variance between the bending stress at each cross-section and the uniform stress state (i.e. the area between the curves in Figure 3) (scalar value ***Y***), (4) iterate on ***X*** until the variance ***Y*** was minimized. The code would then give the drag force profile ***X*** that produced the most uniform stress state along the specimen (***s***), and the total variance in the bending stress (***Y***), which is a quantitative assessment of how efficiently the specimen’s structural tissues (i.e., rind) were allocated. The optimization routine was conducted with several different initial starting points for each specimen (i.e., initial values for F_0_ and f(x)) to ensure the global optimal solution was found as opposed to a local minimum. Tolerances and stopping criteria were set to 1E-6 (first order optimality tolerance), 1E-6 (function tolerance), and 1E-10 (step size tolerance).

**Figure 3:**
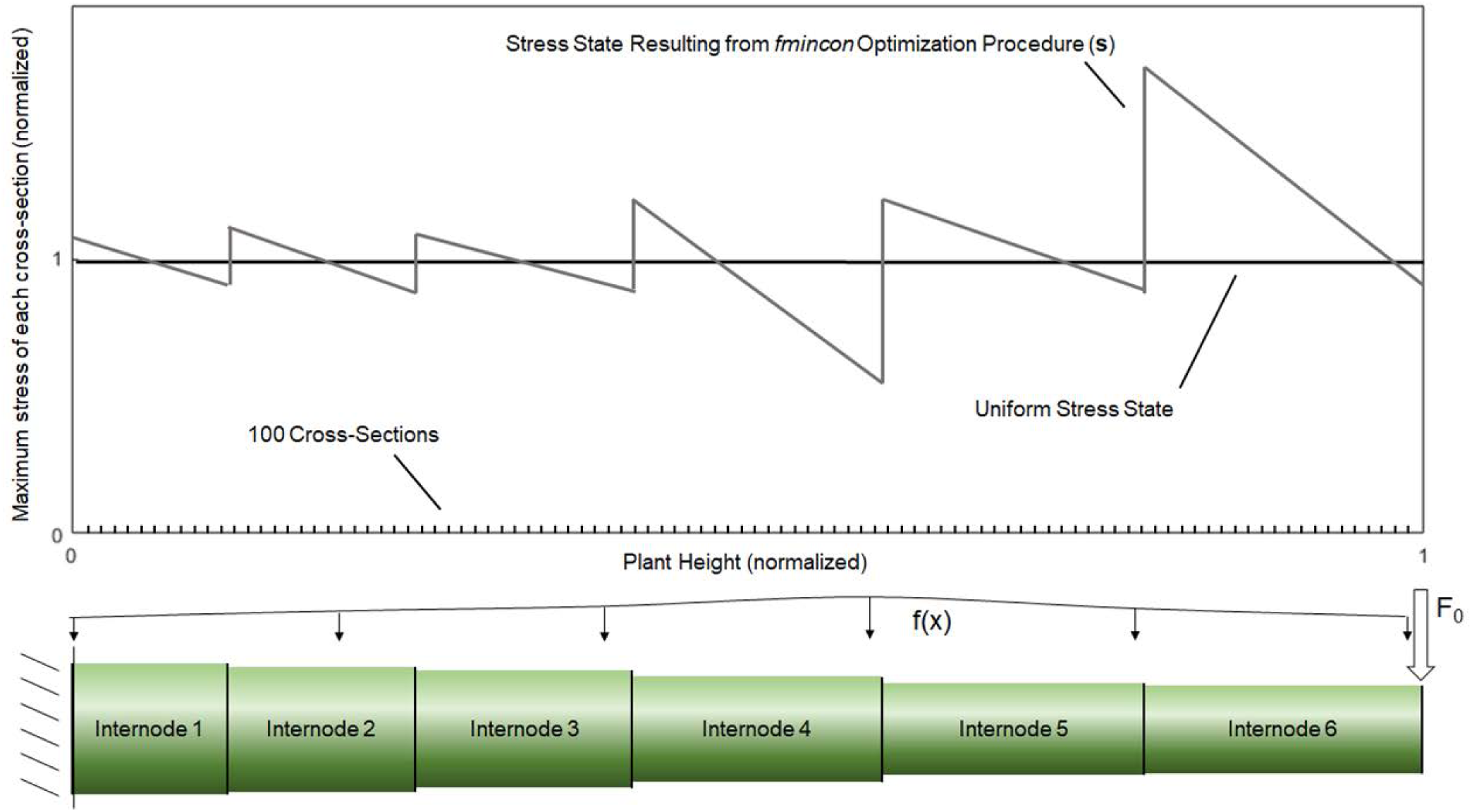
A typical specimen output, showing the stress state resulting from the fmincon optimization procedure, which attempts to create the most uniform possible stress state along the length of the specimen by altering values for F_0_ and f(x) (top); the analytical maize stalk model partitioned into 100 cross-sections along its length (bottom).

## Results

Both the three-point-bending tests and rind penetration tests yielded the expected results, based on previous studies (Robertson *et al*., 2015, 2017; Stubbs *et al*., 2018). The three-point-bending force-deflection responses were linear in nature until failure and demonstrated failure patterns that occur in naturally lodged maize plants (Robertson *et al*., 2015; Stubbs *et al*., 2018). The rind penetration tests gave results characteristic of the protocol. Both test regimes were found to be reliable and repeatable. The bending strength of each specimen, as well as the section modulus, length, major diameter, minor diameter, and rind thickness of each specimen internode is presented in Figure 4.

**Figure 4:**
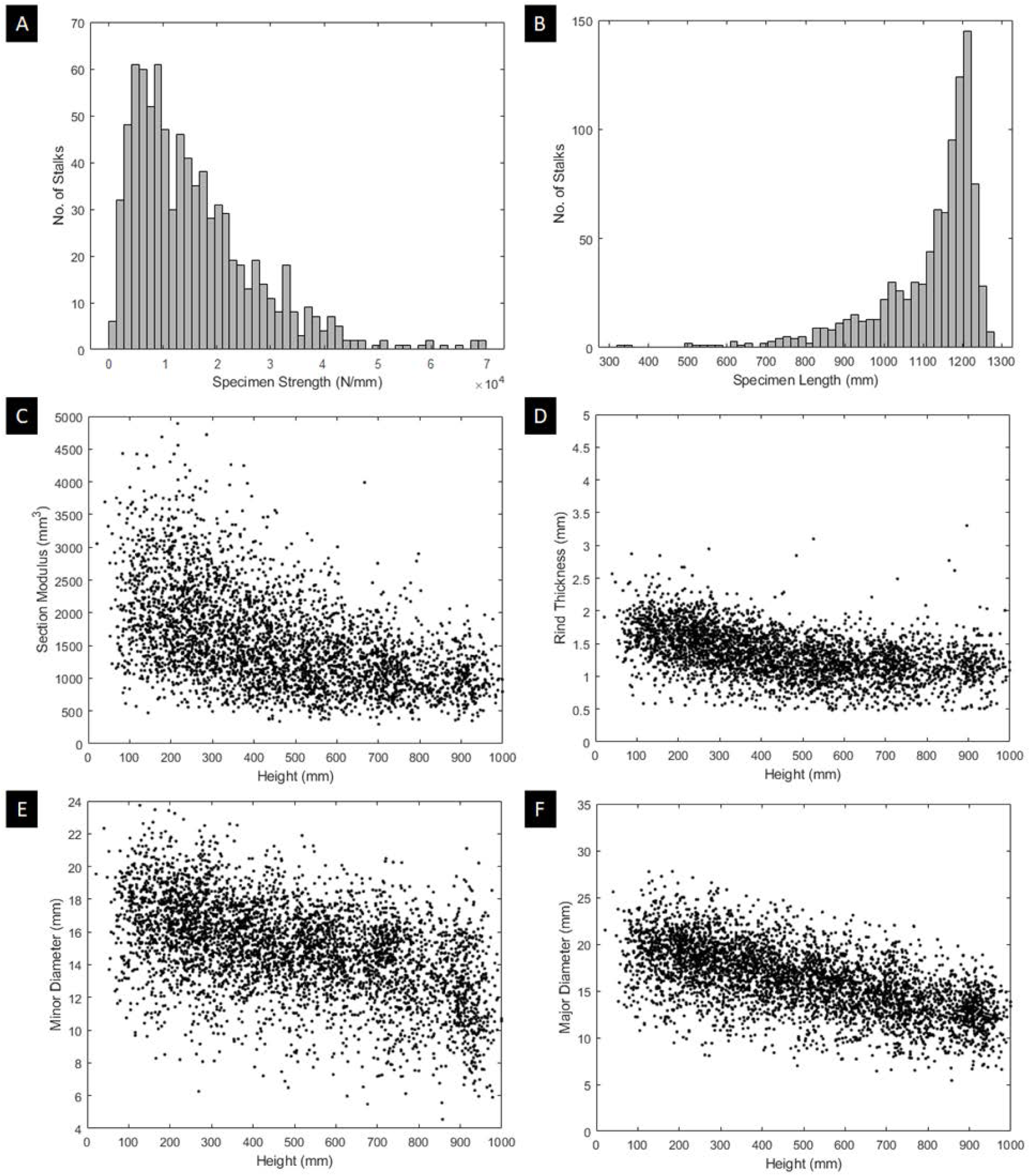
The geometric characterization of all 945 specimens; histograms of the specimen strengths (a) and specimen lengths (b); plots of the section modulus (c), rind thickness (d), minor diameter (e), and major diameter (f) along the lengths of each specimen.

### Structural Efficiency

Section modulus values for each stalk were analyzed to determine structural efficiency (i.e., how structurally efficient the taper of each stalk was). It was found that the median taper of all stalks demonstrated an efficient allocation of structural tissues for probable wind loadings (see Figure 5). However, many internodes fell well outside the range of structural efficiency (i.e., outside of the white area in Figure 5). In particular, 35% of the measured internodes in the study fell within the most efficient range, 38% of measured internodes fell below the blue curve (too little structural tissue), and 27% of the measured internodes fell above the red curve (too much structural tissue).

**Figure 5:**
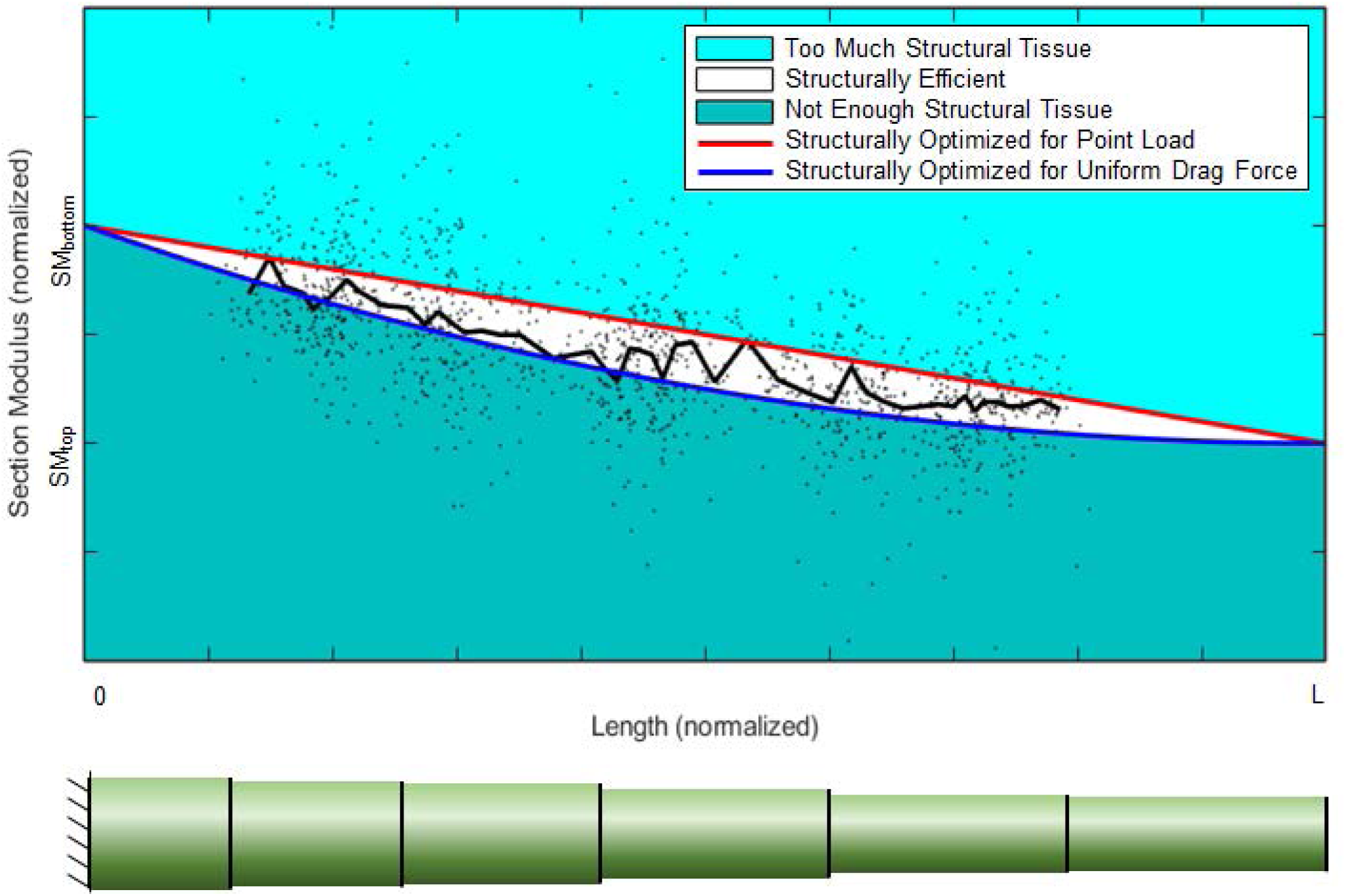
The measured section moduli of the specimens as compared to the region of structural efficiency.

### Optimal Drag Force Profile for Each Stalk

The optimization procedure was performed on all 945 stalks. The *fmincon* procedure successfully determined the drag force profile (***X***) that produced the most uniform state for each specimen. Figure 6 depicts histograms of the resulting stress states of the specimens. In particular, the overall average stress along the length of each specimen (n = 945) and the stress at every cross-section of each specimen (n = 94500) is presented in Figure 6. To enable all specimens to be plotted on the same graph the stress of each specimen / cross-section was normalized to a target stress of 1.00. In other words, a stress state different than a stress of 1.00 represents a suboptimal allocation of structural tissues.

**Figure 6:**
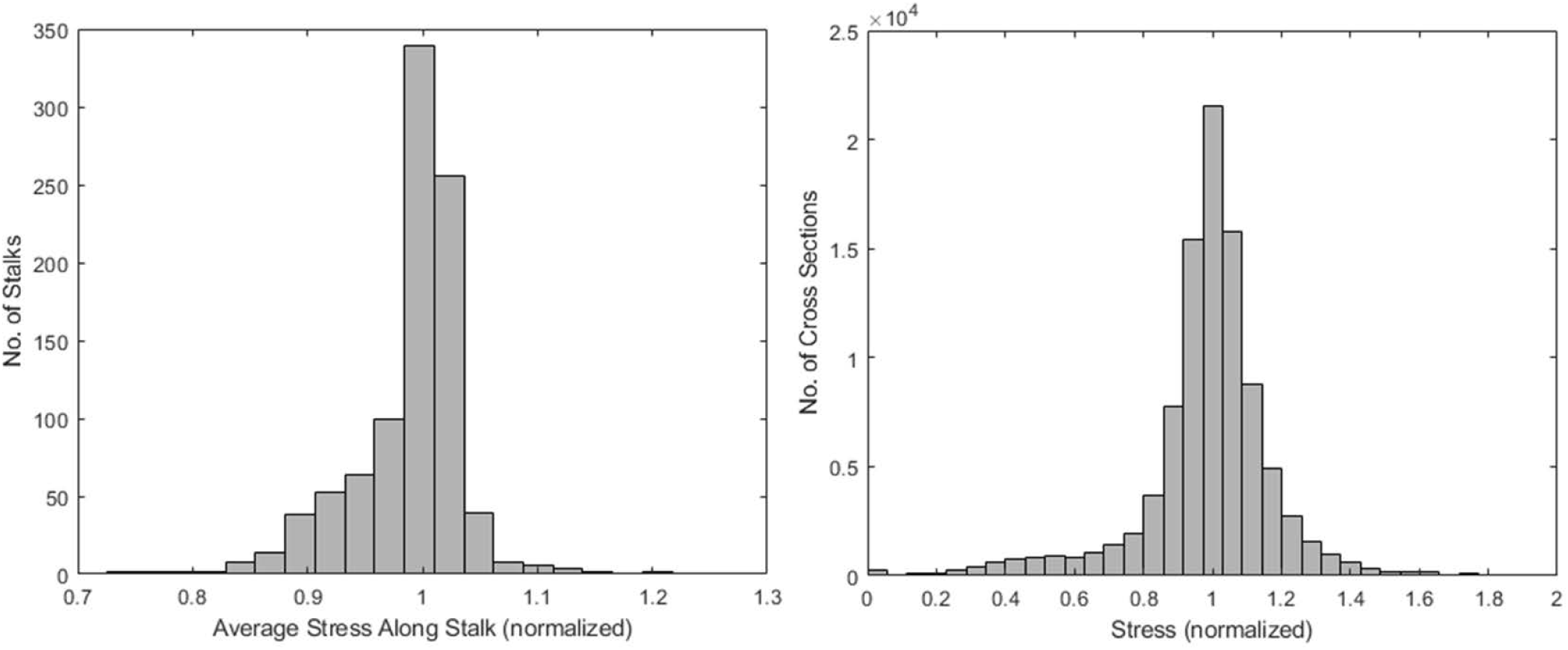
A histogram of the average stress along the length of each of the 945 specimens after optimization, n = 945 (left); a histogram of the average stress at all 100 cross-sections along the length of each of the 945 specimens after optimization, n = 94500 (right). All stress are normalized to a target stress of 1.00. Thus stress states that deviate from 1.00 represent a suboptimal structural allocation of biomass.

Analysis of the drag force profiles for each specimen that would produce the most uniform stress in the specimen revealed that the resolved force F_0_ was far larger than the drag force profile below the ear (see Figure 7). These data imply that the stalks allocate structural tissues for wind loading that primarily occurs above the ear (e.g. the drag force increases exponentially with height). This does not imply that there is no wind below the ear, but that the drag force (determined by the local wind speed, projected stalk and leaf area, and drag coefficient) is much less below the ear as compared to the drag force above the ear. Note this does not imply the bending stresses are lower at the base of the stalk. Bending stresses are determined by forces (i.e., f(x) and F_0_) and moment arms (i.e., distance at which the force is applied). Thus, bending stresses are always higher at the base of the stalk even if the drag force profile is lower at the base of the stalk.

**Figure 7:**
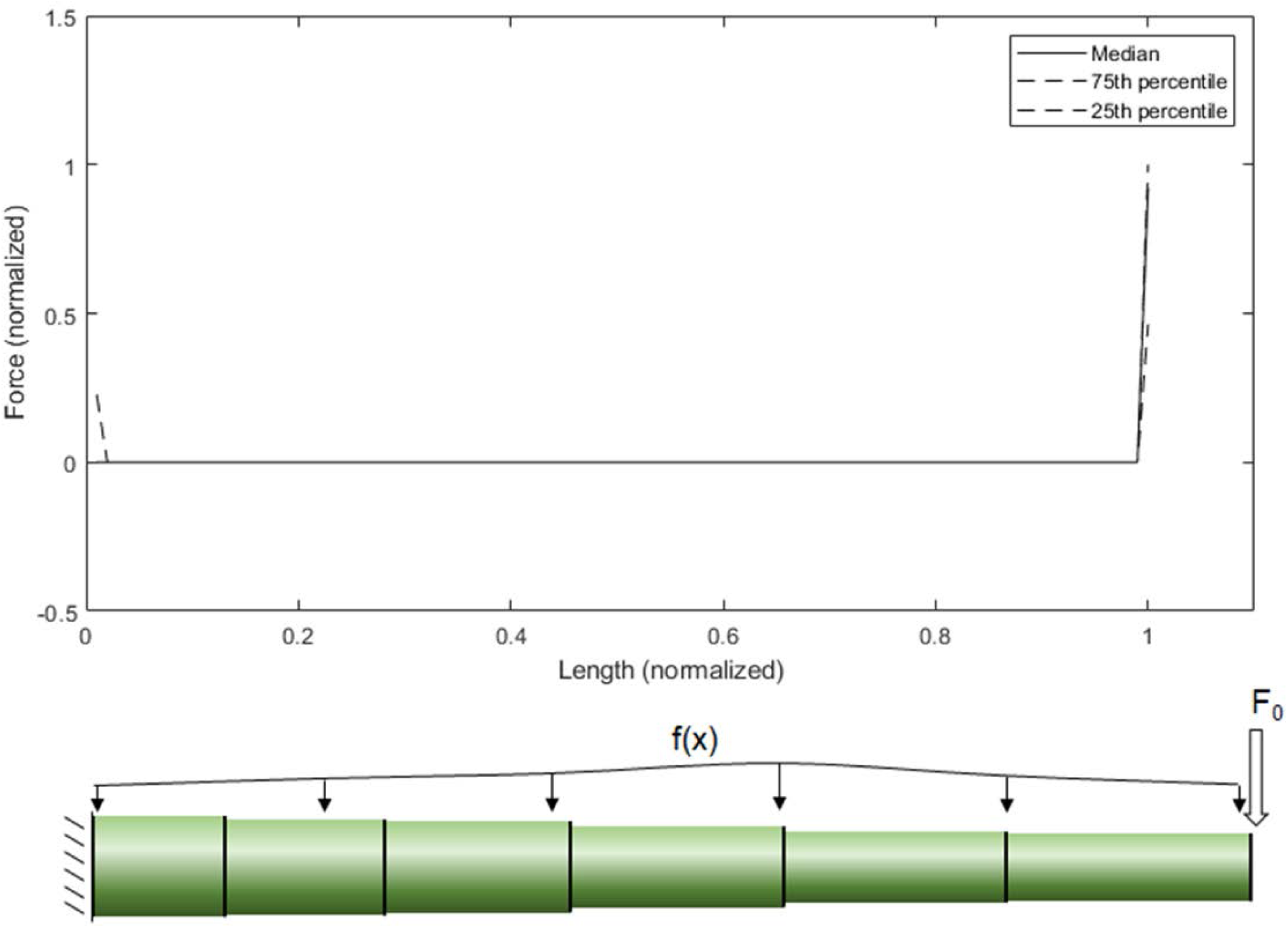
The drag force profile for all 945 specimens (f(x) and F_0_) (top); a loading diagram of f(x) and F_0_ (bottom). The 75th and 25th percentile values are generally line-on-line with the median values along the length of the specimen.

### Are Stronger Stalks More Efficient

To test the hypothesis that stronger plants allocate structural tissues more efficiently (i.e., they produce uniform stresses under probable wind loading scenarios), the bending strength of each stalk was compared to its taper. As discussed previously, the level of structural efficiency can be determined by calculating the area between the curves shown in Figure 3 (***Y***). For example, a ***Y*** value of zero represents a perfectly efficient structure, and the larger the ***Y*** value, the less efficiently the stalk tissues are organized. The data demonstrated that stronger stalks more structurally efficiently (have a lower ***Y*** value) and more consistent (lower variance of ***Y*** values). This was found to be true when comparing individual specimens and when comparing hybrids (see Figure 8).

**Figure 8:**
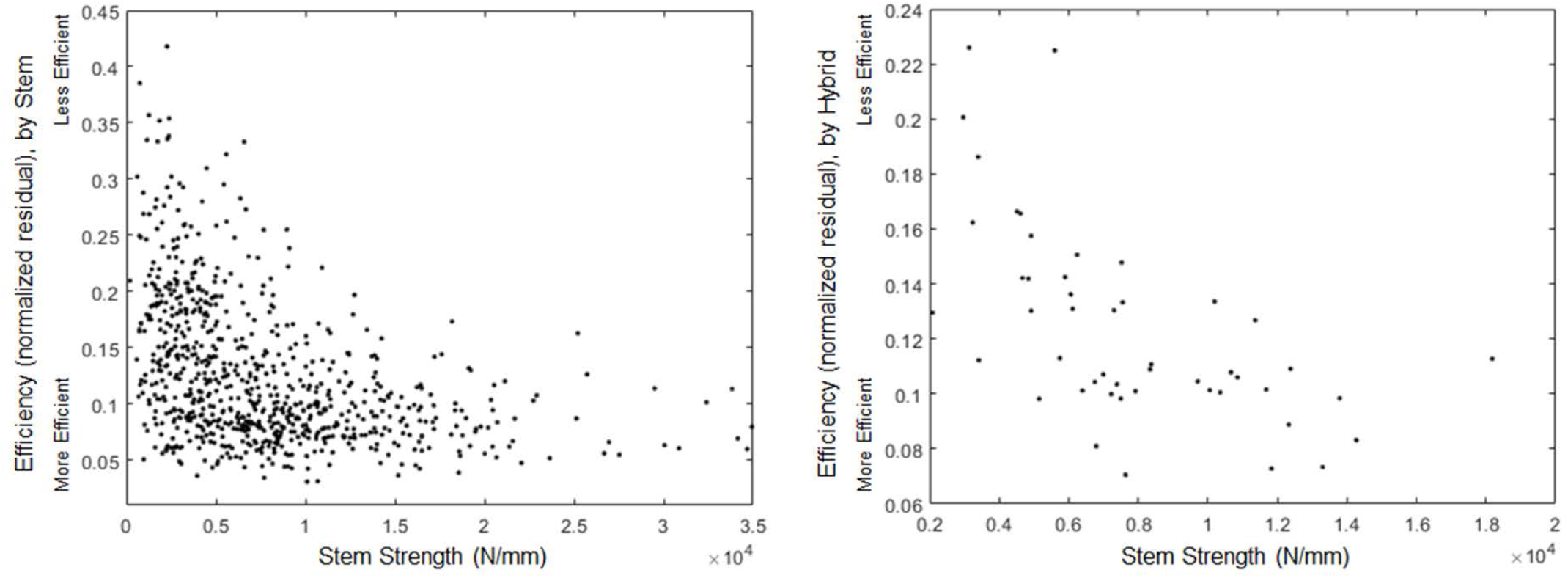
Strength vs. structural efficiency for each specimen (left) and the average of each hybrid (right).

## Discussion

Improving structural efficiency in maize plants could simultaneously enhance yield and lodging resistance. However, there has not been any previous investigations of structural efficiency in maize stalks. Consequently, plant scientists have not directly breed or managed for plants that are structurally optimized in the past. Results from this study suggest that the majority of modern maize hybrids may possess suboptimal stalk structures (see figure 5). In other words, most maize plants utilize bioenergy and structural biomass inefficiently. This reduces the amount of potential biomass and bioenergy available for grain filling (i.e., lowers harvest index) and simultaneously makes stalks more susceptible to stalk lodging. Of the 945,000 cross-sections analyzed in this study 65% were structurally suboptimal with 38% having too much structural tissue and 27% having too little structural tissue.

Further studies are needed to determine the genetic, environmental and management practices that influence stalk taper and structural efficiency in maize stalks. In addition, high throughput phenotyping methods capable of economically measuring stalk rind thickness are needed. Most methods of measuring rind thickness require destructive sectioning and imaging procedures that induce plant fatality and thus prevent measurement of other important crop breeding metrics such as yield. Several nondestructive methods of measuring stalk rind thickness have been developed (e.g., x-ray computed tomography) but these methods are usually limited to laboratory or greenhouse settings and cannot easily be implemented in an agricultural field setting (e.g., (Mairhofer *et al*., 2012; Robertson *et al*., 2017; Seegmiller *et al*., 2020)).

Results showed that lodging resistant hybrids (i.e., those with higher average bending strengths) were more structurally efficient than hybrids that were weaker. The hybrids with higher average bending strengths also displayed less plant to plant variation in structural efficiency. In other words, strong hybrids were more structurally optimized and more consistently optimized than weaker plants (i.e., demonstrated less plant to plant variability). These same findings are true when analyzing individual plants. For example, if the hybrid factor is ignored and each stalk is analyzed as an individual specimen (i.e., no averaging of results across hybrids) the stronger stalks were more structurally efficient than weaker stalks. These results are likely due in part to breeding techniques used in the past. In particular, applied selective breeding pressure based on counts of lodged stalks at harvest time is expected to produce hybrids that are both strong and exhibit minimal plant to plant variance in strength. That is to say that a variety with high average strength but also high standard deviation in strength will have higher lodging rates than a variety with a similar average strength but a lower standard deviation in strength.

In this study an optimization routine was used to determine the wind loading profile that would produce the most uniform mechanical stresses along the length of maize stalks. It was found that maize stalks are structurally optimized for wind loadings that occur primarily above the ear. This is consistent with the authors’ observations in field conditions; although plants at the border of the field may experience loading along the full length of the stalk, the majority of maize plants appear to be primarily subjected to wind loads at or above the ear. The optimization method used in this study was robust and has the potential to be applied to other plants in which the structure of the plant may be predictive of its loading environment. Using optimization methods to infer a plants wind loading environment has several advantages over traditional measurement techniques used to determine wind loads on plants. In particular, it is computationally efficient (as compared to fluid-structure interaction models) and can infer the aggregate loading over time, taking into account the wind profile and fluid-structure interaction between the wind and the plant stalk.

Three-point bending tests are the most commonly employed test to quantify bending strength in plant stalks (Robertson *et al*., 2015; Stubbs *et al*., 2018). However, results from this study highlight several shortcomings of the three-point-bending test approach. In particular, most plant stalks are tapered and researchers typically opt to place the loading anvil from a three-point-bending test at the same anatomical location for each stem specimen in a given study (e.g. the third internode). Thus, the failure location is artificially imposed by the researcher since failure always occurs near the loading anvil, whereas in nature, the failure location is determined by local material weakness and imperfections (i.e., suboptimal allocation of structural tissues). By artificially imposing the failure location the researcher is inducing failure in a cross-section that may have more structural tissue than is optimal in some specimens and less structural tissue than is optimal in other specimens. This confounds comparisons of bending strength among different specimens in a given study, as the measured bending strength could vary substantially for any given specimen depending on the structural optimality of the failed cross-section. A better approach is to apply bending loads that replicate natural loading patterns. Such loading conditions produce natural failure types and failure patterns in plant specimens (i.e., failure occurs at the cross-section with the least optimal allocation of structural tissue). Several devices have been recently developed which accomplish this task (Grafius and Brown, 1954; Berry *et al*., 2003; Guo *et al*., 2018, 2019; Erndwein *et al*., 2019; Heuschele *et al*., 2019). In particular, they utilize the natural anchoring of the maize roots and apply a point load to a cross-section near the ear (very similar to the loading profile shown in Figure 7). Thus these devices simulate the loading conditions experienced by plants in their natural environment and consequently produce natural failure types and patterns (Cook *et al*., 2019). These devices are therefore expected to provide more distinguishing power than three-point-bending test methods.

### Limitations

The primary limitation of the current study is that the rind of the stalk was assumed to be a homogeneous, isotropic, linear elastic material subjected to pure bending. The inclusion of heterogeneity, anisotropy, non-linear material properties, and the addition of the pith material could change the behavior of the analyzed stalks. However, previous research has shown the inclusion of these effects to be small and in many cases insignificant. The authors do not believe inclusion of such effects would change the overall conclusions of this paper (Von Forell *et al*., 2015; Al-Zube *et al*., 2018).

This study is deliberately limited in its scope to structural efficiency. Other abiotic and biotic considerations can affect stalk morphology / anatomy and should be considered in future studies. In addition, this study utilized three-point-bending test to measure the bending strength of stem specimens. However, as mentioned in the discussion section these tests are less than ideal. Unfortunately, at the time the study was conducted by the authors we were not fully aware of the limitations of three-point-bending test. In particular, we did not expect to find that the majority of maize stalk cross-sections are structurally suboptimal. Future studies are warranted which utilize in-field phenotyping devices (Erndwein *et al*., 2019) to assess structural efficiency and its relations to stalk strength, harvest index, etc.

## Conclusions

The morphology of physiological mature maize stalks was characterized, and the loading environments that result in the most uniform maximum stresses along the length of maize stalk were investigated. It was found that maize stalks are morphologically organized to resist wind loading that occurs primarily above the ear. It was also found that plants with higher bending strengths were more structurally efficient than weaker plants. However, even strong plants allocated structural tissues in a suboptimal manner. There exists much room for improvement in the area of structural optimization of maize stalks. These findings are relevant to crop management and breeding studies seeking to improve stalk lodging resistance.

## Authors’ Contributions

All authors were fully involved in the study and preparation of the manuscript. The material within has not been and will not be submitted for publication elsewhere.

## Acknowledgements

This work was funded in part by the National Science Foundation (Award #1826715) and by the United States Department of Agriculture - NIFA (#2016-67012-2381). Any opinions, findings, conclusions, or recommendations are those of the author(s) and do not necessarily reflect the view of the funding bodies.

